# Probiotic-guided CAR-T cells for universal solid tumor targeting

**DOI:** 10.1101/2021.10.10.463366

**Authors:** Rosa L. Vincent, Candice R. Gurbatri, Andrew Redenti, Courtney Coker, Nicholas Arpaia, Tal Danino

## Abstract

Synthetic biology enables the engineering of interactions between living medicines to overcome the specific limitations of any singular therapy. One major challenge of tumor-antigen targeting therapies like chimeric antigen receptor (CAR)-T cells is the identification of targetable antigens that are specifically and uniformly expressed on heterogenous solid tumors. In contrast, certain species of bacteria selectively colonize immune-privileged tumor cores and can be readily engineered as antigen-independent platforms for therapeutic delivery. Bridging these approaches, we develop a platform of probiotic-guided CAR-T cells (ProCARs), in which T cells are engineered to sense synthetic antigens (SA) that are produced and released by tumor-colonizing probiotic bacteria. We demonstrate increased CAR-T cell activation and tumor-cell lysis when SAs anchor to components of the extracellular matrix. Moreover, we show that ProCARs are intratumorally activated by probiotically-delivered SAs, receive further stimulation from bacterial TLR agonists, and are safe and effective in multiple xenograft models. This approach repurposes tumor-colonizing bacteria as beacons that guide the activity of engineered T cells, and in turn builds the foundation for communities of living medicines.

## Main Text

While there has been significant success in the use of chimeric antigen receptor (CAR)-T cells for hematological malignancies, translation to solid tumors has been greatly limited. The central challenge of antigen-targeted cell therapies is intrinsically tied to the expression patterns of the antigen itself – making the identification of an optimal target one of the most critical factors in the development of new CARs for solid tumors^1–3^. Few tumor-associated antigens (TAAs) identified on solid tumors are tumor-specific, and carry a high risk of fatal on-target, off-tumor toxicity due to cross-reactivity against vital tissues^4–6^. Moreover, if a safe target can be identified, TAAs are often heterogeneously expressed and selection pressures from targeted therapies ultimately lead to antigen-loss and tumor escape^7,8^. Emerging strategies to address the antigen bottleneck have focused on improving CAR-T cells with additional genetic circuitry^9–11^, modulatory proteins^12–15^, and combinations with nanoparticles and oncolytic viruses^16–18^.

Where CAR-T cells require significant engineering to target and infiltrate solid tumors, bacteria naturally colonize immune-privileged tumor cores and preferentially grow within hypoxic and necrotic tumor microenvironment (TME)^19^. Indeed, a multitude of patient studies have now shown that different tumor types host unique tumor microbiomes^20–22^. Taking advantage of these inherent properties, several groups have established an array of synthetic gene circuits to engineer a new class of prokaryotic, cell therapy^23,24^. These approaches have utilized engineered bacteria as intratumoral bioreactors that continually produce a range of payloads, resulting in tumor regression and mitigation of systemic side effects^25–28^. Importantly, clinical trials with engineered bacteria have thus far reported minimal toxicities in patients with solid tumors, though have yet to demonstrate considerable clinical efficacy across a broad range of indications^29–31^.

Here, we bridge the complementarity of these two cell therapies by creating a platform of probiotic-guided CAR-T cells (ProCARs), whereby T cells are engineered to sense and respond to synthetic targets that are released by tumor-colonizing, probiotic bacteria. Ultimately, this approach leverages the antigen-independence of tumor-seeking microbes to create a combined cell therapy platform for universal solid tumor targeting.

## Results

### ProCAR-T cells are designed to sense and respond to synthetic antigens produced by intratumoral bacteria

Toward the goal of creating a tumor-antigen independent ProCAR system, we engineered synthetic antigens (SA) that can be readily produced by tumor-colonizing bacteria, and exclusively released within the tumor core based on a quorum-regulated genetic circuit. Specifically, we chose a well-characterized probiotic strain, *E. coli* Nissle 1917 (EcN), equipped with a genomically-integrated synchronized lysis circuit (synchronized lysing integrated circuit, SLIC) for intratumoral SA delivery. Importantly, we have demonstrated that the SLIC system cyclically permits bacterial growth to reach a critical population density within the tumor before triggering synchronized lysis events that act to release genetically-encoded payloads within TME^32^.

To ensure orthogonality to healthy human tissue, SAs were designed from two components: (1) a homodimer of sfGFP (D117C)^33^, previously shown to mediate CAR-T cell responses to dimeric forms of soluble factors^34^, and (2) a known heparin binding domain from placenta growth factor-2 (PIGF-2_123-144_)^35^ linked to sfGFP with a flexible glycine-serine linker (PIGF), which can anchor to the high levels of collagens and fibronectins found within the dense extracellular matrix (ECM) of solid tumors^36^. We hypothesized that an ECM-binding SA would benefit the ProCAR system twofold: (1) by limiting SA diffusion beyond ECM-dense tumor margins, thereby enhancing the safety of the system, and (2) facilitating CAR polarization akin to conventional CAR and TCR signaling^37^, resulting in greater antitumor activity (**Fig. 1A**). We separately cloned both SA designs downstream of polyhistidine-tags (His-tag) for protein purification (**Fig. S1A, B**), and under a constitutive *tac* promoter for high protein expression *in vivo* from an Axe/Txe stabilized plasmid, as previously described (**Fig. S2A, B**)^38^. We then confirmed efficient collagen binding of PIGF-linked sfGFP following purification of His-tagged SA (**Fig. S1C**).

**Figure 1.**
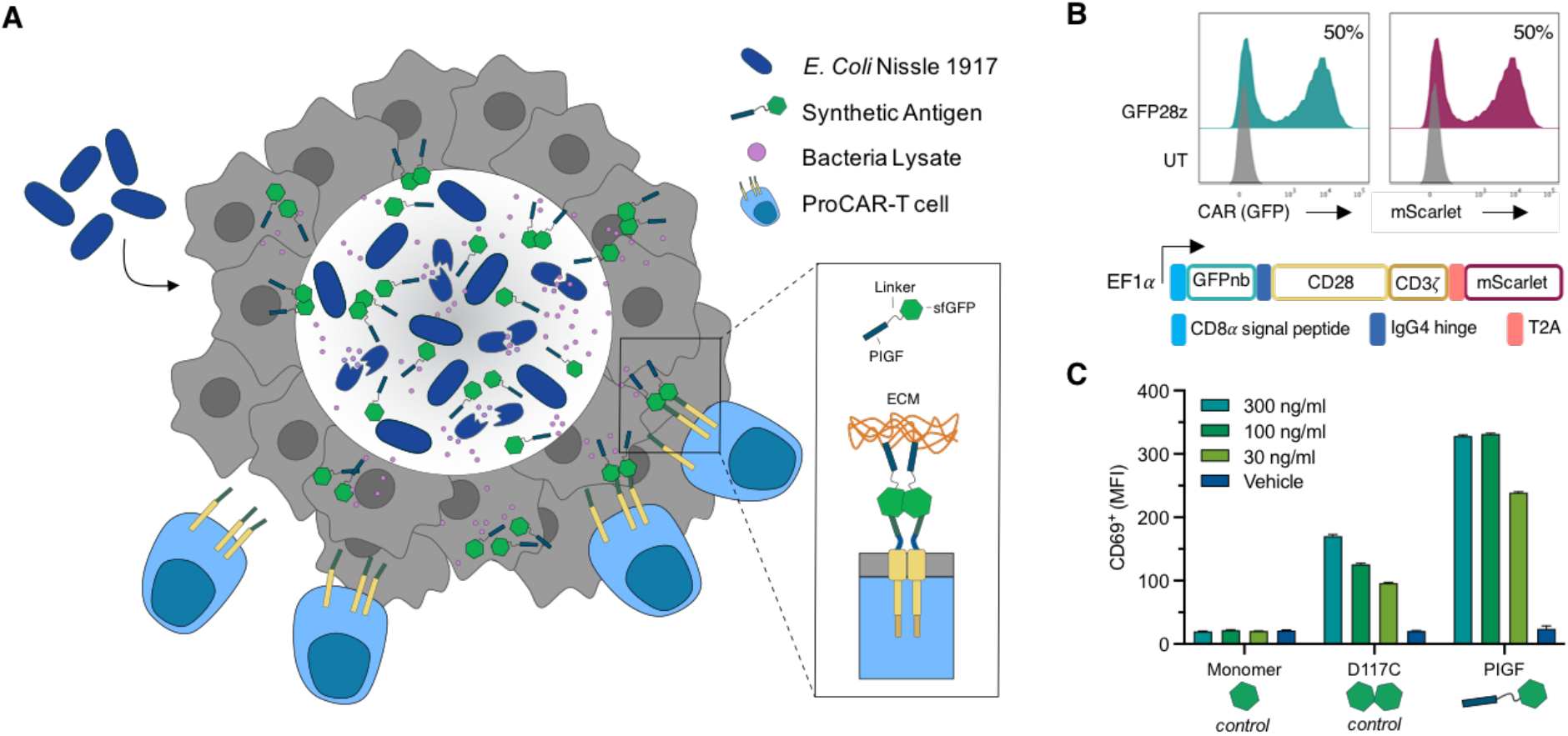
Probiotic-guided CAR-T cells (ProCARs) are programmed to sense and respond to synthetic antigens (SA) released by tumor-colonizing probiotics. (**A**) Schematic of the ProCAR system whereby engineered *E. coli* Nissle 1917 specifically colonize the solid tumor core and release synthetic antigens (SA) through quorum-regulated lysis. SAs are designed with sfGFP and PIGF sequences to anchor to components of the extracellular matrix (ECM) and locally activate GFP-specific ProCAR-T cells within the TME. (**B**) Representative flow cytometry histograms demonstrating GFP-CAR (GFP28z) surface expression and binding to purified monomeric-sfGFP (left) in primary human T cells together with the co-expression of the fluorescent marker, mScarlet (right). GFP28z contains an extracellular sfGFP-binding nanobody linked to CD28 and CD3ζ intracellular domains through a short IgG4 hinge and CD28 transmembrane domain, and co-expression of mScarlet is achieved through separation by a T2A element. (**C**) Flow cytometric quantification of CD69 surface expression on GFP28z exposed to collagen-bound PIGF relative to monomeric and dimeric (D117C) sfGFP controls.

To complete the GFP-based system we composed an SA-specific CAR based on an sfGFP-binding nanobody sequence^39^ linked to CD28 and CD3ζ intracellular signaling domains by a short IgG4 hinge. We then constructed a lentiviral vector to co-express the CAR gene and a fluorescent mScarlet reporter from the EF1α promoter, separated by a self-cleaving T2A element. With this, we were able to monitor transduction efficiency of human T cells by mScarlet expression, and confirm surface expression and target specificity through CAR receptor binding to purified, monomeric sfGFP (**Fig. 1B**). With the SA-system components in place, we sought to assess the extent of CAR-T cell activation elicited by each purified sfGFP variant. Here we found that GFP CAR-T cells (GFP28z) were strongly activated by collagen-bound PIGF, moderately activated by dimeric D117C, and remained unchanged by exposure to monomeric sfGFP (**Fig. 1C, S1D**).

### GFP-directed CAR-T cells mediate killing of target cells in response to collagen-bound sfGFP

To further assess this pattern of activation, we next interrogated the surface expression of additional activation markers and intracellular cytokine production by flow cytometry. As anticipated, we observed the highest surface levels of CD25 on GFP28z incubated with collagen-bound PIGF, and lower expression in response to soluble D117C (**Fig. 2A**). This trend was again mirrored in the intracellular levels of Th1 proinflammatory cytokines detected after 16 hr of co-incubation, with GFP28z producing the highest levels of IFNγ, IL-2, and TNFα, in response to collagen-bound PIGF (**Fig. S3A**). Similarly, these cells displayed an increased frequency of polyfunctional CD8^+^ T cells producing both IFNγ and TNFα (**Fig. 2B**). Stronger PIGF-mediated activation did not appear to drive higher rates of T cell expansion than exposure to D117C; with GFP28z cells expanding to a peak of 8-fold when stimulated with either SA (**Fig. 2C**). This may be attributable to the comparable levels of IL-2 detected in cell culture supernatants by 24 hours of exposure to PIGF or D117C (**Fig. S3B**).

**Figure 2.**
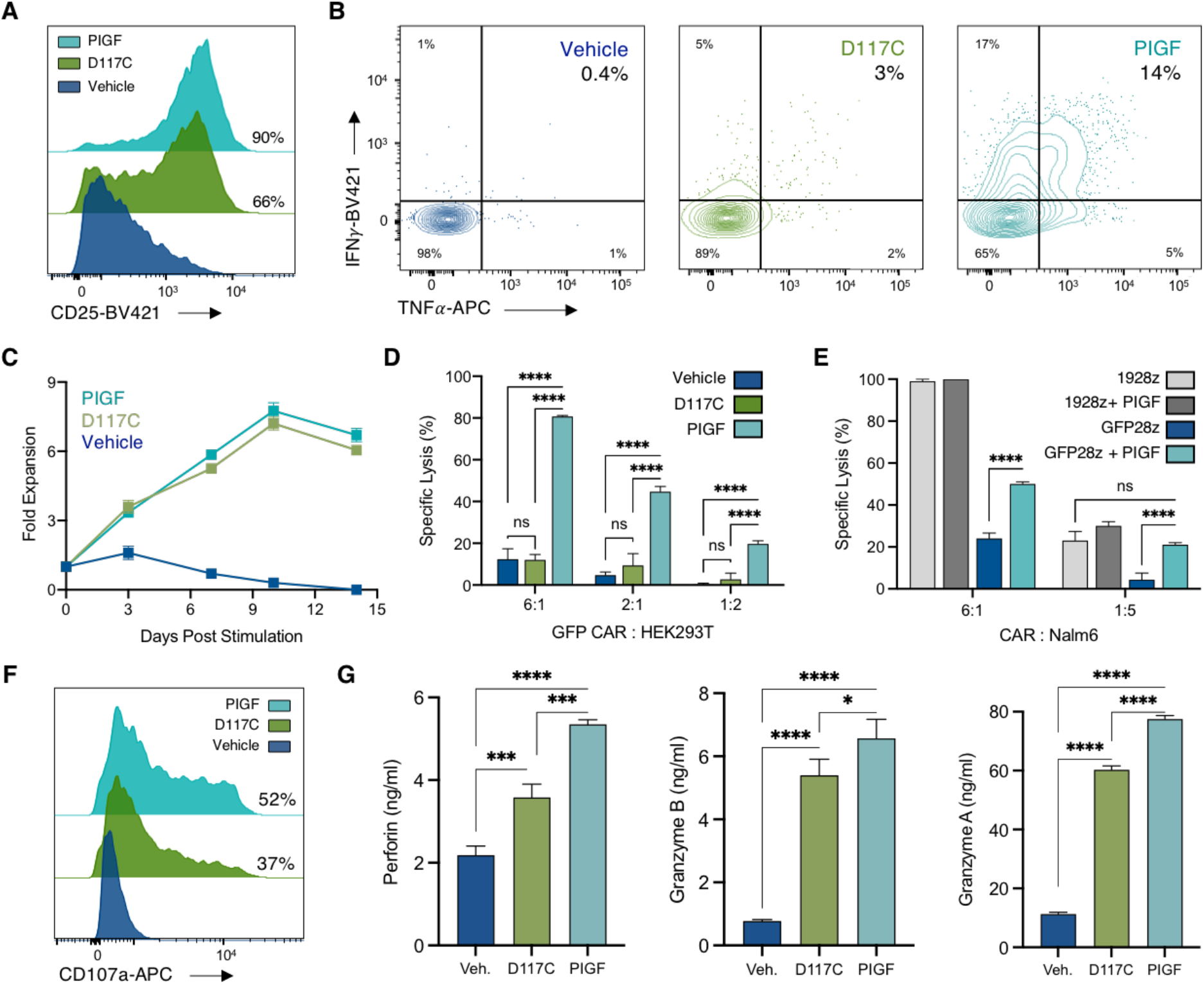
GFP-directed CAR-T cells (GFP28z) activate, and mediate killing of target cells in response to collagen-bound sfGFP. (**A**) Representative flow cytometry histograms of CD25 surface expression on GFP28z following 16hr incubation with 0.1ug/ml of purified SA variants (D117C or PIGF), or a PBS vehicle, on collagen coated plates. (**B**) Representative flow cytometry plots of intracellular levels of IFNγ and TNFα in CD8^+^ GFP28z, treated as in (A). (**C**) SA-driven fold expansion of GFP28z. On day 14 post-activation, GFP28z T cells were washed of IL-2 and exposed to 0.1ug/ml of D117C, PIGF, or PBS vehicle. Cells were counted every 3-4 days for 14 days post stimulation. (**D**,**E**) Overnight killing assays against *ff*Luc^+^ HEK293T (**D**) and *ff*Luc^+^ Nalm6 (**E**) target cells at defined effector to target ratios. CAR-T cells were co-cultured with target cells on collagen coated plates +/−0.1 ug/ml sfGFP-PIGF for 20 hours before lysis and addition of luciferin. Luminescence (RLU) was detected with a Tecan plate reader. Specific lysis (%) was determined by normalizing RLU to co-cultures with untransduced T cells. (**F**) Representative flow cytometry histograms of CAR^+^CD8^+^ T cell degranulation 16-hours post incubation with D117C or PIGF as in (A), and staining for CD107a surface expression. (**G**) Quantification of cytokine levels in cell culture supernatants from GFP28z exposed to PBS (Veh.), D117C, or PIGF for 24hr. Error bars represent s.d. of biological replicates, * p<0.05, ** p<0.01,**** p<0.0001 2-way ANOVA (D,E) or 1-way ANOVA (G), Holm-Sidak multiple comparison correction. ns, not significant; RLU, relative luminescence units.

While previous studies did not observe target cell lysis in response to soluble ligands^34,40^, we hypothesized that the PIGF design would augment lysis of adherent cell lines by co-localizing with the pseudo-ECM of collagen coated plates. Indeed, we observed minimal targeted lysis of *ff*Luc^+^ HEK293T cells when GFP28z cells were supplied with PBS (Vehicle), or D117C, at any effector to target (E:T) ratio; conversely, GFP28z cells incubated with the PIGF modification were able to drive target cell lysis in 80% of HEK293T cells at a high E:T ratio of 6:1, and 20% of cells at a lower E:T ratio of 1:2 (**Fig. 2D**). GFP28z also showed a dose-dependent response to PIGF, with specific lysis of HEK293T cells observed at doses as low as 1.5 ng/ml (**Fig. S4A**). We next tested the effect of PIGF on GFP28z against Nalm6 leukemia cells, a commonly used target of CD19-directed CARs. As anticipated, PIGF had no effect on the potency of CD19 CAR-T cells (1928z). However, GFP28z achieved up to 50% lysis of the suspension cells when supplied with PIGF, with target cell death comparable to that of 1928z at the lower, 1:5, E:T ratio (**Fig. 2E**). We next demonstrated specific lysis of HUH7 hepatoma cells, MCF7 breast cancer cells, and MDA-MB-468 TNBC cells, shown relative to levels of target cell death achieved by GPC3- and ICAM1-targeted CARs, respectively (**Fig. S4B**). Together suggesting PIGF provides a synthetic target that promotes tumor killing across different cell types.

Although high levels of IFNγ secreted by activated T cells can mediate bystander killing of antigen-negative tumor cells^41,42^, we additionally hypothesized that PIGF binding in close proximity to the target cell would facilitate more direct mechanisms of T cell cytotoxicity. Thus, we measured the surface expression of CD107a - a membrane-bound molecule commonly used as a proxy for cytotoxic degranulation, and found higher expression on GFP28z treated with collagen-bound PIGF than cells exposed to D117C (**Fig. 2F**). Moreover, we were able to detect increased levels of perforin and granzymes A/B in the supernatant of GFP28z supplied with either SA, with the highest levels secreted in response to bound antigen (**Fig. 2G**). Taken together, these results suggest that CAR-T cells can mediate effective, tumor cell type–agnostic killing in response to synthetic targets.

### PIGF-based ProCAR system mediates localized anti-tumor activity in a subcutaneous xenograft model of leukemia

Motivated by the cytotoxicity observed *in vitro*, we sought to characterize the full effects of the ProCAR platform *in vivo*, first utilizing NSG mice bearing subcutaneous Nalm6 tumors. We have previously shown that EcN-SLIC (Pro^X^) strains are able to exclusively grow to a critical population density within the tumor core, synchronously lyse, and cyclically release therapeutic payloads every 48-72 hr following a single intratumoral (I.T) injection^32^. Therefore, we chose to treat tumors with a single I.T. injection of 1×10^5^ CFU of Pro^X^ strains either producing the D117C (Pro^D117C^) or PIGF (Pro^PIGF^) SAs, or an empty control (Pro^−^) 48-72 hr before mice received an I.T. injection of 2.5×10^6^ GFP28z cells, or a PBS control (**Fig. 3A**). Here, GFP28z in combination with Pro^D117C^ had no effect on tumor growth, with tumors growing at a similar rate to tumors receiving control Pro^−^ strains alone, or in combination with GFP28z (**Fig. 3B, S5A**). However, Pro^PIGF^ strains were able to mediate a potent antitumor response from GFP28z, leading to significantly slowed tumor growth of Nalm6 tumors and an increased survival benefit (**Fig. 3B, 3C**). This trend was also observed in mice bearing subcutaneous Raji tumors (**Fig. S5B**).

**Figure 3.**
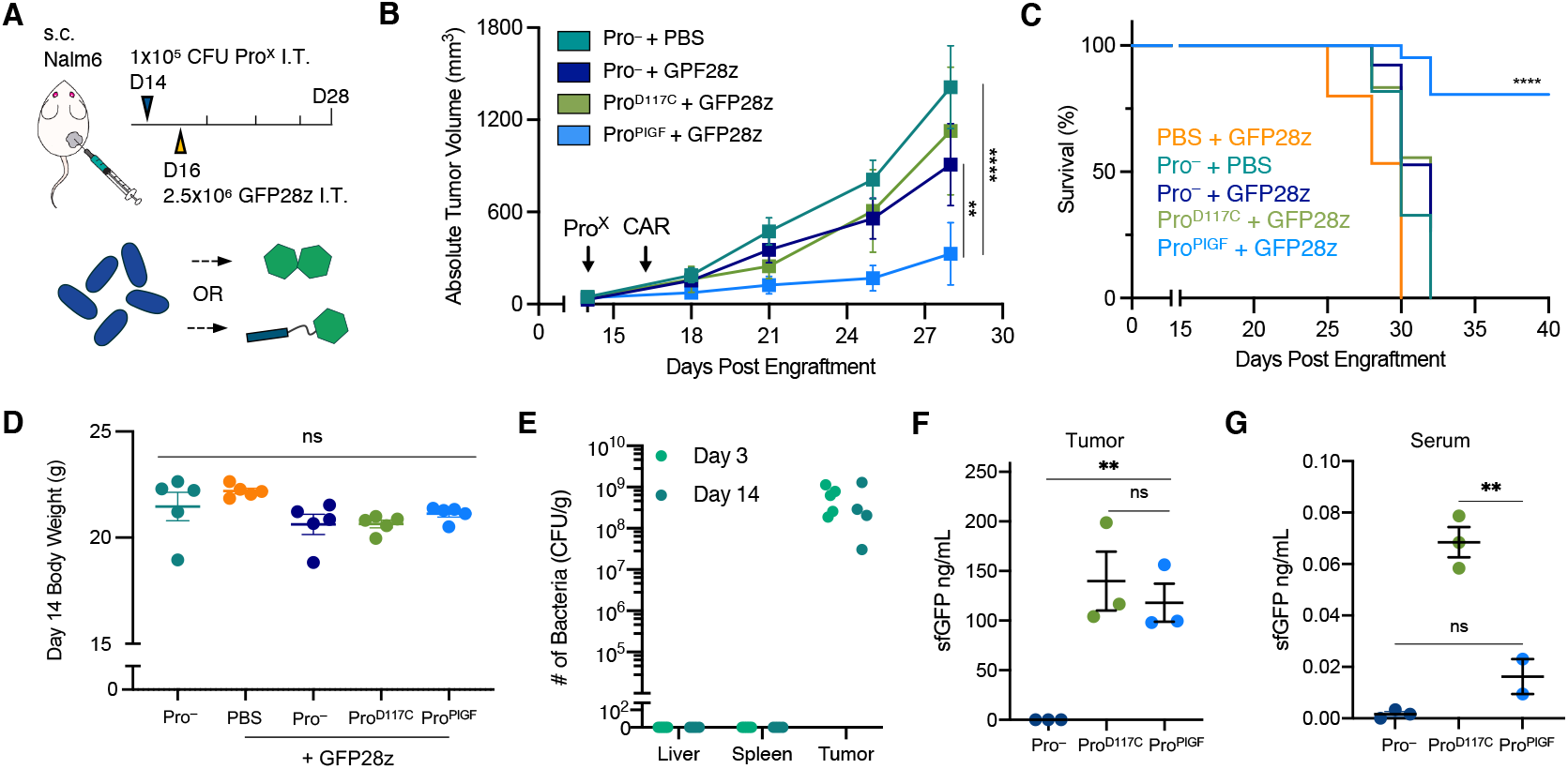
PIGF-based ProCAR system drives localized anti-tumor activity of GFP28z in a subcutaneous xenograft model of human leukemia. (**A**-**D**) Nalm6 cells (5×10^5^) were implanted subcutaneously (s.c.) into the hind flank of NSG mice. When tumor volumes reached ~100mm^3^, mice were intratumorally (I.T.) injected with 1×10^5^ CFU of engineered probiotic strains (Pro^X^) producing D117C (Pro^D117C^) and PIGF (Pro^PIGF^) SA variants, or empty Pro controls (Pro^−^). 2.5×10^6^ GFP28z ProCAR-T cells were then I.T. delivered 48-hours post bacterial injection, with tumor growth monitored by caliper measurements and body weight recorded every 3-4 days (n>4 per group). Mean absolute tumor trajectories (**B**), survival curves (**C**), and mouse body weights at day 14 post bacteria treatment (**D**) are shown. (**E**) Biodistribution of Pro^X^ found in liver, spleen, and tumor samples calculated as colony-forming units (CFU) per gram of tumor. At day 3 (n=5) and day 14 (n=4) post bacteria treatment, tumor, liver, and spleen were homogenized and plated on LB agar plates containing the appropriate antibiotics for bacteria colony quantification. (**F**,**G**) ELISA quantification of sfGFP levels from tumor homogenates (**F**) and serum (**G**) taken 14 days post treatment. Error bars represent s.e.m. of biological replicates, *p<0.05, **p<0.01 ***p<0.001, ****p<0.0001, 2-way ANOVA (B) or 1-way ANOVA (D,F,G), with Holm-Sidak multiple comparison correction. Survival curve (C) ****p<0.0001, log-rank test.

While the safety of bacteria cancer therapies has been tested across multiple syngeneic models^27–29,32,43^, we sought to investigate the safety of the ProCAR system in severely immunocompromised, NSG mice. As a proxy for mouse health and system-tolerance, we monitored mouse body weight from the start of bacteria treatment and observed no significant weight loss in mice treated with GFP28z alone, Pro^−^ alone, or any combination of the two cell therapies (**Fig. 3D**). Moreover, we observed tumor-restricted growth of bioluminescent bacteria *in vivo* (**Fig. S6A**) and did not detect bacteria outside of tumor homogenates on day 3 and 14 post treatment through *ex vivo* assessment of healthy tissues (**Fig. 3E**). Encouragingly, bacteria isolated from Pro^PIGF^-treated tumors at day 14 additionally demonstrated SA-plasmid maintenance (**Fig. S6B**).

To assess the tumor-retention of probiotically-delivered SA variants, we quantified the level of detectable SA in tumor homogenates and matched serum samples from mice treated with Pro^−^, Pro^D117C^, and Pro^PIGF^ strains through GFP-specific ELISA. We detected SA levels at, or higher than, the concentration used in our *in vitro* assays in tumors treated with Pro^D117C^ and Pro^PIGF^ strains, and found no difference in the intratumoral levels of either (**Fig. 3F**). Therefore, the difference in therapeutic efficacy observed between the two groups was likely not the result of differing SA abundance, and attributable to the ability of PIGF to promote target cell killing where D117C does not. As anticipated, higher concentrations of sfGFP were detected in the serum of mice treated with Pro^D117C^, suggesting that ECM-bound PIGF promotes tumor retention and reduces leakage into systemic circulation (**Fig. 3G**). Taken together, these findings motivated the utility of PIGF as the SA design of the ProCAR platform.

### Engineered strains of *E. coli* Nissle 1917 enhance CAR-T cell effector function

The heart of the ProCAR platform is an unlikely team of two cell types whose direct interaction is not well characterized. However, it has been shown that activated T cells upregulate TLR4 and TLR5 expression^44,45^ – of which, LPS and flagellin from EcN are respective agonists – therefore, we hypothesized that intratumoral bacterial lysate may serve as an adjuvant to enhance ProCAR-T cell activity. To test this *in vitro*, we monitored the surface expression of CD69, CD25, and CD107a, on GFP28z cells exposed to media alone, EcN lysate, PIGF alone, or the combination of PIGF and EcN lysate. GFP28z demonstrated significantly elevated levels of all three markers in response to EcN lysate alone, with the combination of lysate and collagen-bound PIGF stimulating the highest levels (**Fig. 4A**). Furthermore, CD8^+^ T cells exposed to EcN lysate displayed an effector-differentiated phenotype, with terminally differentiated effector populations (T_eff_, CD45RO^−^CD62L^−^) expanding in both untransduced and GFP28z T cells (**Fig. 4B, S7**). GFP28z exposed to the combination displayed the strongest enrichment of T_eff_ populations and furthest reduction in central memory populations (T_cm_, CD45RO^+^CD62L^−^), while stem cell memory populations (T_scm_, CD45RO^−^CD62L^+^) were maintained. The synergistic effect of PIGF in combination with EcN lysate was again mirrored in levels of Granzyme B and pro-inflammatory cytokines, GM-CSF, IFNγ, IL-2, and TNFα, detected in cell culture supernatants (**Fig. 4C**).

**Figure 4.**
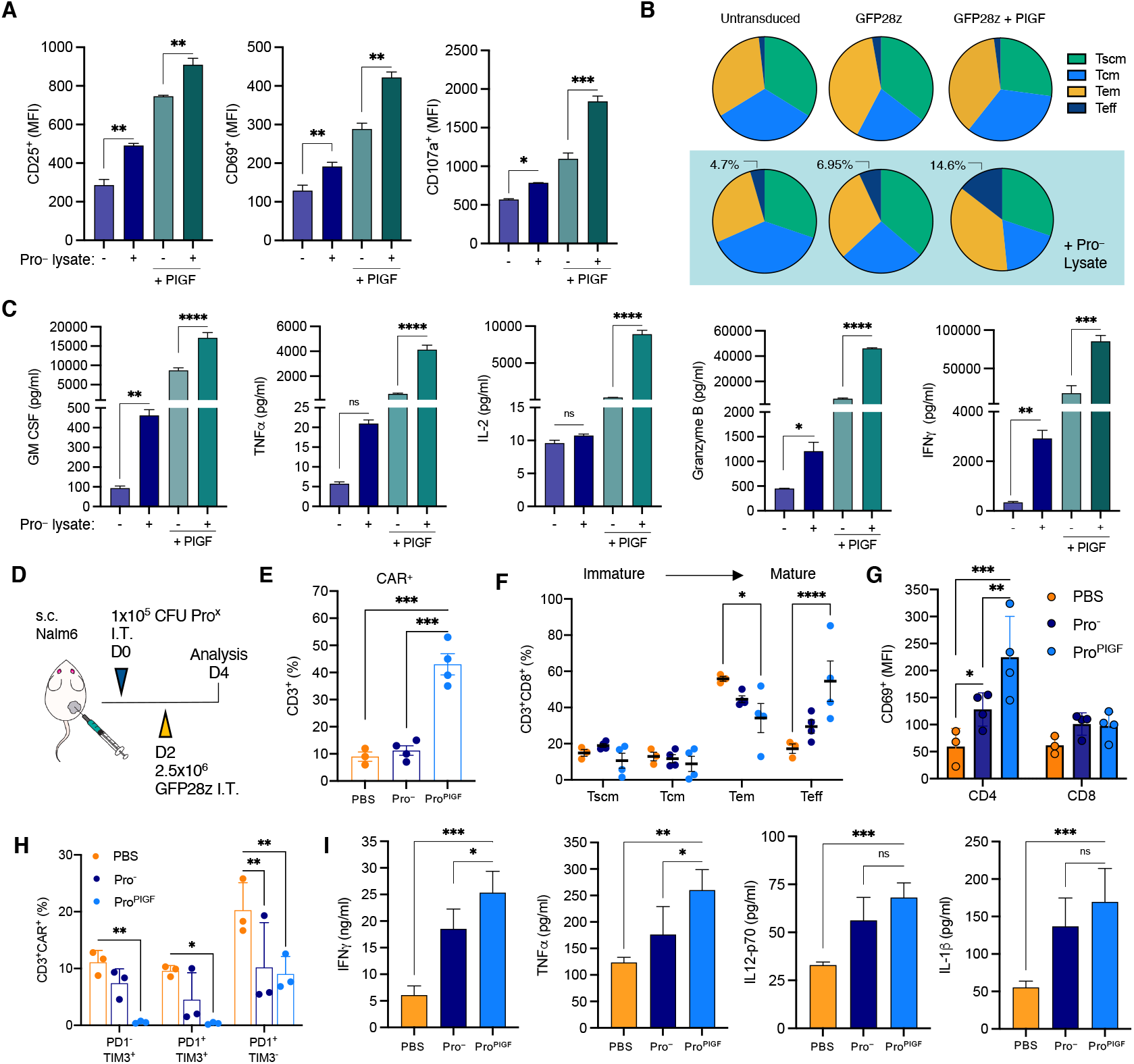
Engineered Pro^X^ strains of *E. coli* Nissle 1917 (EcN) enhance T cell effector function. (**A**) Quantification of flow cytometric analysis assessing surface expression of CD25, CD69, and CD107a in response to EcN lysate +/−PIGF. GFP28z cells were plated on collagen-coated plates and exposed to lysate at a final OD of 1, +/−0.1 ng/ml purified PIGF. (**B**) Phenotype of T-cells exposed to stimulants as in (A), for 48hr. Pie charts representing the clockwise differentiation of CD8^+^ T cell populations from stem cell memory (T_scm_) CD62L^+^CD45RO^−^, to central memory (T_cm_) CD62L^+^CD45RO^+^, to effector memory (T_em_) CD62L^−^CD45RO^+^, and to terminal effector (T_eff_) CD62L^−^CD45RO^−^ cells. (**C**) Quantification of cytokine levels detected in cell culture supernatants from GFP28z treated as in (A). (**D**-**I**) Nalm6 cells (5×10^5^) were implanted subcutaneously into the hind flank of NSG mice. When tumor volumes reached ~100 mm^3^, mice were I.T. injected with either PBS or 1×10^5^ CFU of Pro^PIGF^, or Pro^−^ controls. On day 2 post Pro^X^ injection, all groups received an I.T. injection of 2.5×10^6^ GFP28z ProCAR-T cells. Tumors were harvested and homogenized on day 4 for analysis. (**E**) Frequency of intratumoral CAR^+^ T cells as determined by flow cytometric detection of GFP-bound hCD45^+^CD3^+^ cells. (**F**) Frequency of intratumoral hCD45^+^CD3^+^CD8^+^ T cell memory and effector populations determined by CD62L and CD45RO expression patterns as in (B). (**G**) Flow cytometric quantification of CD69 surface expression on intratumoral hCD45^+^CD3^+^ CD8^+^ or CD4^+^ cells in each treatment group. (**H**) Frequency of intratumoral hCD45^+^CD3^+^CAR^+^ cells displaying single positive PD1^−^TIM3^+^, double positive PD1^+^TIM3^+^, or single positive PD1^+^TIM3^−^ surface expression within each treatment group. (**I**) Luminex quantification of intratumoral cytokine levels. Error bars represent s.d. of biological replicates, *p<0.05, **p<0.01 ***p<0.001, ****p<0.0001, 1-way ANOVA, with Holm-Sidak multiple comparison correction.

To study the effects of PIGF produced by live bacteria, we interrogated the phenotype of GFP28z isolated from Nalm6 tumors treated with PBS, Pro^−^, or Pro^PIGF^. Tumors received an I.T. injection of Pro^X^ strains before receiving a second I.T. injection of GFP28z on day 2 post treatment, tumors were then homogenized and prepared for flow cytometry on day 4 (**Fig. 4D**). Here, tumors from all three groups were found to contain comparable levels of human CD45^+^CD3^+^ cells (**Fig. S8A**). However, we observed a significant increase in the frequency of CD45^+^CD3^+^CAR^+^ cells in tumors treated with Pro^PIGF^, suggesting CAR^+^ populations were specifically expanding in response to locally-released SA from tumor-colonizing bacteria (**Fig. 4E**). Moreover, CD8^+^ GFP28z T cells from Pro^PIGF^-treated tumors were significantly enriched for terminally differentiated T_eff_ populations, while cells from Pro^−^-treated tumors displayed a modest trend toward T_eff_ differentiation (**Fig. 4F**).

As a measure of activation, CD4^+^ GFP28z cells displayed significantly increased CD69 expression in response to Pro^−^ and Pro^PIGF^ strains *in vivo* (**Fig. 4G**), though CD25 expression was found only to increase in response to Pro^PIGF^ (**Fig. S8B**). Interestingly, the exhaustion phenotype of GFP28z appeared inversely correlated with exposure to either of the Pro^X^ strains, likely attributable to the beneficial effects of MyD88-costimulation following TLR activation^46^. We observed the highest frequency of PD-1^−^TIM-3^+^, PD-1^+^TIM-3^+^, and PD-1^+^TIM-3^−^ cells in the PBS control, while GFP28z from Pro^PIGF^-treated tumors were found to be absent of the exhaustion marker TIM-3 entirely, and T cells from Pro^−^-treated tumors displayed an intermediate phenotype (**Fig. 4H**). Indeed, cytokine profiling from tumors treated as above revealed a similar pattern. Tumors treated with either Pro^X^ strains were found to contain significantly increased levels of multiple human, pro-inflammatory cytokines, IFNγ, TNFα, IL12-p70, and IL-1β, relative to PBS groups, with the highest levels of IFNγ and TNFα detected in Pro^PIGF^-treated tumors (**Fig. 4I**). Together, these observations highlight the dual functionality of the Pro^PIGF^ strain by providing both synthetic CAR targets and natural TLR stimulants that reshape the TME for enhanced CAR-T cell effector function.

### The ProCAR system produces a durable anti-tumor response in a subcutaneous xenograft model of triple negative breast cancer

We next sought to test the ProCAR platform in mice bearing subcutaneous MDA-MB-468 TNBC tumors to establish TAA-independent efficacy in an additional solid tumor model. As before, tumors were treated with an I.T. injection of Pro^X^ strains, or PBS, two days prior to receiving an I.T. injection of either PBS, GFP28z, or an ICAM1-directed CAR (ICAM28z, **Fig. 5A**). Interestingly, the combination of Pro^PIGF^ and GFP28z demonstrated enhanced antitumor efficacy relative to ICAM28z, despite high ICAM1 expression on MDA-MB-468 cells^47^. While Pro^PIGF^ again mediated the strongest antitumor activity of GFP28z, the therapeutic effects of TLR stimulation alone were evident in the combination treatment of GFP28z with Pro^−^ (**Fig. 5B, S9A**). However, by day 51 post engraftment (day 25 post treatment) the therapeutic benefit of Pro^PIGF^ and GFP28z appeared reduced, and by day 55 we observed only a small difference between the weights of excised tumors from groups treated with GFP28z in combination with PBS, Pro^−^, or Pro^PIGF^ (**Fig. S9B**). Upon phenotypic interrogation of the excised tumors, we observed higher counts of human CD45^+^CD3^+^ cells (**Fig. 5C**) and significantly increased counts of CAR^+^ cells in Pro^PIGF^-treated tumors (**Fig. 5D**). As anticipated, the majority of these cells were classed as CD62L^−^CD45RO^−^ T_effs_ across all three groups (**Fig. S9C**), and displayed high levels of the exhaustion markers LAG-3, TIM-3, and PD-1 (**Fig. 5E**) – suggestive of the T cell dysfunction commonly observed in the immunosuppressive TME^1^. Encouragingly, *ex vivo* assessment of Pro^−^ and Pro^PIGF^ bacterial isolates from the same tumors revealed preserved functionality of quorum-based lysis and PIGF production (**Fig. S9D**). Following these observations, we sought to prolong the antitumor activity of the ProCAR system by fixing treatment at one dose of the Pro^X^ strains and increasing the frequency of T cell treatment to two doses, spaced two weeks apart (**Fig. 5F**). With this, the Pro^PIGF^ and GFP28z combination was able to achieve a durable antitumor response, with no tumor growth observed 70 days post engraftment (**Fig. 5G, S10**).

**Figure 5.**
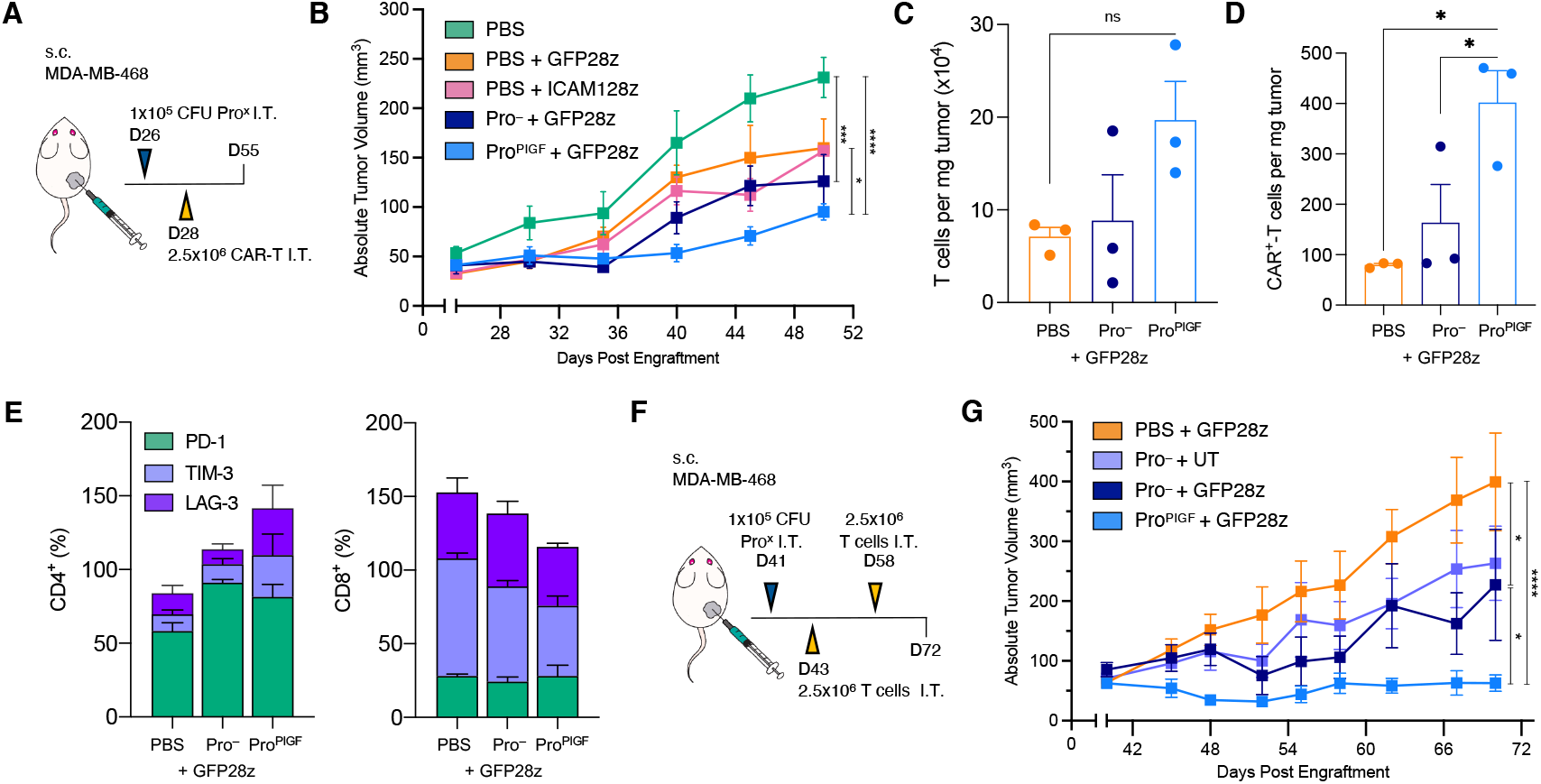
ProCAR system drives durable anti-tumor effects in a subcutaneous xenograft model of triple negative breast cancer (TNBC). (**A**-**E**) MDA-MB-468 cells (5×10^5^) were implanted subcutaneously (s.c.) into the hind flank of NSG mice. When tumor volumes reached ~50mm^3^, mice were I.T. injected with either PBS or 1×10^5^ CFU of Pro^PIGF^ or empty Pro^−^ controls. Two days post injection, mice received a single I.T. injection of either PBS, 2.5×10^6^ GFP28z ProCAR-T cells, or 2.5×10^6^ ICAM1-specific CAR-T cells (ICAM28z), with tumor growth monitored by caliper measurements every 3-4 days (n>4 per group). Mean tumor trajectories (**B**) are shown. (**C-E**) Tumors treated with PBS, Pro^−^, and Pro^PIGF^ in combination with GFP28z were harvested and prepared for flow cytometry on day 30 post bacteria treatment (D55 post tumor engraftment). (**C**) Absolute counts of hCD45^+^CD3^+^ cells per mg of tumor. (**D**) Absolute counts of hCD45^+^CD3^+^CAR^+^ cells per mg of tumor. (**E**) Frequency of intratumoral CD4^+^ and CD8^+^ T cells displaying exhaustion markers, LAG-3, TIM-3, or PD-1, within each treatment group. (**F-G**) S.c. MDA-MB-468 tumors were established in NSG mice prior to I.T. injection with PBS, Pro^−^, or Pro^PIGF^ as in (A). Mice then received an initial I.T. injection of 2.5×10^6^ untransduced (UT), or 2.5×10^6^ GFP28z ProCAR-T cells two days post bacteria treatment, followed by a second I.T. dose of UT or GFP28z 14 days later. Tumor growth was monitored by caliper measurements every 3-4 days (n>4 per group). Mean tumor trajectories are shown (**G**) Error bars represent s.e.m. of biological replicates, *p<0.05, **p<0.01 ***p<0.001, ****p<0.0001, 2-way (B, G) or 1-way ANOVA (C,D), with Holm-Sidak multiple comparison correction.

## Discussion

Here, we have presented a new approach to engineering interactions between living therapies, where tumor-colonizing probiotics have been repurposed as beacons that guide and direct the activity of engineered T cells. We have shown that by fusing synthetic CAR targets to a heparin binding domain we can achieve TAA-independent target cell death, and by harnessing the tumor-restricted growth of *E. coli*, we can release these targets directly within the TME to achieve tumor-agnostic efficacy in genetically distinct models of human leukemia and TNBC. Our findings underscore the potential of the ProCAR platform to potentially surpass the critical roadblock of identifying suitable CAR targets by providing an antigen that is orthogonal to both healthy tissue and tumor genetics.

Notably, even the gold standard CD19 CAR faces antigen-dependent issues, the loss of which has become a frequent cause of patient relapse^7^, and off-tumor expression on brain mural cells has been linked to reports of dangerous neurotoxicity^5^. This antigen problem plays a larger role for solid cancers, and approaches to overcome these issues have primarily relied on incorporating complex genetic circuitry to afford greater control over TAA recognition. Strategies targeting more than one antigen circumvent issues of tumor escape^48,49^, however, the challenge of finding a single suitable target has limited this approach for most solid tumors.

Tumor pattern recognition and Boolean-gated logic circuits provide elegant solutions to the issue of off-tumor toxicity; however, they involve complex T cell engineering and insertion of multiple transgenes that potentially limit translation^11^. Flexible approaches to build ‘universal CARs’, where the antigen recognition domain is provided by separate I.V. infusion, enables exchanging of antigen specificity during treatment without the need for additional transgenes^12,50^. Additionally, CARs secreting BiTEs against broadly expressed targets have been effective in preclinical models of heterogenous tumors^14^. While these strategies represent significant advancements, these approaches do not aim to provide both a ‘universal CAR,’ and a tumor-independent ‘universal antigen.’

With this in mind, several groups have looked to nanoparticles and oncolytic viruses (OVs) as alternate platforms to deliver T cell targets directly to the tumor^16–18^. Recently, TNBC and colorectal tumors were cleared using a combination of a CD19 CAR and an OV delivering the CD19 ectodomain to tumor cells^17^. The authors highlight consequential B cell aplasia as a mechanism to avoid antibody responses against the OVs, however, the approach could be extended to the delivery of orthogonal CAR targets at the cost of repeat dosing. Nonetheless, the use of bacteria in the ProCAR system offers a partner organism that is amenable to complex engineering^51^, including the incorporation of additional circuits to restrict growth and immunogenicity in ways that facilitate safe systemic delivery and repeat dosing^52–55^. Moreover, the current study has highlighted natural inflammatory properties of probiotics that serve to enhance T cell effector function in an otherwise immunosuppressive-TME. Future iterations will involve probiotic strain engineering to balance the consequential effector differentiation observed here with mechanisms to enhance T cell homeostasis for more durable responses.

Importantly, the ProCAR system presents a valuable opportunity to modulate the TME beyond T cell engineering. Future work will leverage the modularity of tumor-homing probiotics to remodel the TME to actively recruit and sustain engineered T cells through the delivery of additional payloads including chemokines^56^, PD-1 inhibitors^57^ and critical cytokine support^58^. Moreover, with the lack of treatment options for patients with metastatic disease, it will become important to explore systemic delivery of the ProCAR platform. Intravenously delivered *E. coli* has been shown to safely colonize tumors as small as 1 mm^3^ in multiple mouse models^19^, however, systemic delivery has yet to be tested in severely immunocompromised systems. Beyond this, the use of NSG mice to assess the combination of probiotics with human T cells has also limited the assessment of potential inflammatory side effects including cytokine release syndrome^59^.

Nonetheless, the current study has demonstrated tumor-restricted growth of bacteria in an immunocompromised context and established the ProCAR system as a well-tolerated and effective therapeutic platform across multiple models. Altogether, combining the advantages of tumor-homing bacteria and CAR-T cells provides a new strategy for tumor recognition, and in turn, builds the foundation for engineered communities of living therapies.

## Methods

### Study Design

The overall purpose of this study was to provide proof-of-concept work demonstrating the effective combination engineered probiotics with compatibly-engineered CAR-T cells by harnessing the most beneficial characteristics of two distinct cellular therapies. We tested an ECM-binding SA against a soluble SA using several tumor models, techniques, and approaches. These employed three xenogeneic models and *in vitro* characterization of CAR-T cell function across five distinct target cell lines. Each experiment was performed multiple times with T cells isolated from multiple healthy human donors.

### DNA constructs

The GFP specific ProCAR construct was synthesized (IDT) based on a previously reported amino acid sequence for a GFP-binding nanobody^39^, the CD19 CAR was constructed from the FMC63 antibody, the GPC3 CAR was constructed from the GC33 antibody, and the ICAM1 CAR was constructed by using the ligand binding domain from LFA1 (LFA1_129-318_)^60^. All antigen-recognition domains were fused to an IgG4 hinge and CD28 transmembrane and intracellular costimulatory domains in tandem with a CD3ζ signaling domain, and were linked to mScarlet by a T2A peptide sequence. CAR genes were cloned into a modified pHR_SFFV lentiviral transgene expression vector (Addgene plasmid #79121, deposited by Wendell Lim) with the EF1α promoter inserted in place of SFFV using EcoRI and NotI restriction digest and In-Fusion cloning (Takara Bio). Synthetic antigens (SAs) were constructed by synthesizing two geneblocks (IDT) encoding a *tac* promoter and *E. coli*-optimized genes for D117C and PIGF with N-terminal His-tags. Specifically, D117C was constructed using the sfGFP coding sequence with a cysteine substitution at position D117^33^ and PIGF was constructed by linking sfGFP to the PIGF_123-144_ peptide sequence (RRRPKGRGKRRREKQRPTDCHL)^36^ with a flexible glycine-serine linker (G_4_S_1_) at the C-terminus. All bacterial payloads were cloned into an Axe/Txe stabilized p246-AT plasmid^38^ using Gibson assembly (NEB) methods.

### Purification of synthetic antigens (SA)

D117C and PIGF SA variants were cloned into an inducible expression vector and transformed into eNiCo21(DE3) E. *coli*. Transformants were grown at 37°C to an OD_600_ of ~0.9 and induced with 1 mM IPTG for 16 hr at 30°C. Cells were then centrifuged for 10 min at 4000 rpm and resuspended in lysis buffer (50 mM NaH_2_PO_4_, 300 mM NaCl, pH 8.0) for sonication. Lysates were spun for 30 min, following which the supernatant was loaded onto Ni-NTA (Qiagen) resin, washed in wash buffer (35 mM imidazole), and eluted in 250 mM imidazole for collection. The elutions were dialyzed in PBS using regenerated cellulose dialysis tubing (3500 Da MWCO) and then filtered through a 0.2-μm filter. 488 nm absorbance was used to determine the concentration, and diluted to a final of 1 mg/mL in PBS ready for use, aliquots were stored at −80°C.

### Primary human T cell isolation and culture

Primary human T cells were isolated by negative selection for CD3^+^ populations (STEMCELL technologies, Easy Sep) from anonymous healthy human donor blood collected by leukopheresis and purchased from STEMCELL technologies. T cells were cryopreserved in CryoStor10 (STEMCELL technologies). After thawing, T cells were cultured in human T cell medium (hTcm) consisting of X-VIVO 15 (Lonza) and 5% Human AB serum (Gemini) supplemented with 50 units/mL IL-2 every 2 days for all experiments (Miltenyi Biotec).

### Generation of human CAR-T cells

Pantropic VSV-G pseudotyped lentivirus was generated by transfection of HEK293T (ATCC, CRL-11268) with a pHR’SIN:CSW transfer vector and psPAX.2 and pMD2.G packaging plasmids using Lipofectamine 3000 (Invitrogen). 24 hr post transfection, T cells were thawed and activated with anti-CD3/CD28 DYNAL™ Dynabeads™ (Gibco) at a 1:2 cell:bead ratio. At 48 hr post transfection and day 1 post T cell activation, viral supernatants were harvested and added to T cells at an MOI of 1.5-2 with 0.8 μg/mL of polybrene (MilliporeSigma). T cells were exposed to the virus for 24 hr before removal of polybrene and addition of fresh hTcm. Dynabeads were removed from T cell cultures at day 6 post activation and T cells were transferred to GRex24 (Wilson Wolf), or GRex6, plates for expansion until day 13-14, at which point were considered rested and ready for use in assays or cryopreservation.

### Pro^X^ strain generation and administration

D117C, PIGF, and CXCL9 expression vectors were transformed into electrocompetent EcN-SLIC^32^ strains and cultured in LB media with 50 μg/ml kanamycin with 0.2% glucose, in a 37°C shaking incubator. For therapeutic preparation, Pro^X^ strains were grown overnight in LB media containing appropriate antibiotics and 0.2% glucose. The overnight culture was sub-cultured at a 1:100 dilution in 50 mL of fresh media with antibiotics and glucose and grown to an OD_600_ of ~ 0.05, preventing bacteria from reaching quorum. Cells were centrifuged at 3000 rcf and washed 3 times with sterile ice-cold PBS. Pro^X^ strains were then diluted to a final concentration of 5×10^6^ CFU/mL in cold PBS, 40 μL of each strain was then injected intratumorally.

### Cell lines

Jurkat Clone E6-1 cells were purchased from ATCC (TIB-152) and lentivirally transduced to stably express either the GFP or CD19 CARs and cultured in RPMI-1640 (Gibco) supplemented with 10% fetal bovine serum (FBS, Gibco) and 1% penicillin/streptomycin (Gibco). Adherent target cell lines were purchased from ATCC unless otherwise stated and cultured in DMEM/F12 supplemented with 10% FBS and 1% penicillin/streptomycin: HEK293T cells (CRL-11268), MCF7 breast cancer cells (HTB-22), MDA-MB-468 triple negative breast cancer (HTB-132), and the HUH7 hepatoma cells were a gift from R. Schwabe (Columbia University). All target cell lines were transduced to stably express firefly luciferase (*ff*Luc), *ff*Luc^+^ Nalm6 acute lymphoblastic leukemia cells were a gift from M. Sadelain (Memorial Sloan Kettering Cancer Center) and cultured in RMPI-1640 supplemented as above.

### Animal models

All animal experiments were approved by the Institutional Animal Care and Use Committee (Columbia University, protocols AC-AAAN8002 and AC-AAAZ4470). Mice were blindly randomized into treatment groups. Animal experiments were performed on 6-8-week old female *NOD.Cg-Prkdc*^*scid*^ *Il2rg*^*tm1Wjl*^/*SzJ* (NSG) mice (Jackson Laboratory) with subcutaneous hind-flank tumors from implanted human acute lymphoblastic leukemia cells (Nalm6), or Burkitts lymphoma cells (Raji, ATCC CCL-86), or triple negative breast cancer cells (MDA-MB-468). All tumor cells were prepared for implantation at a concentration of 5×10^6^ cells/mL in PBS and matrigel (Corning) at a 1:1 ratio. Tumors were grown to an average volume of approximately 100 mm^3^ before receiving an intratumoral injection of bacteria as described above. Two days later mice received an intratumoral injection of 2.5×10^6^ CAR^+^ T cells (5×10^6^ total T cells) in 50 μL of PBS. Tumor volume was calculated by measuring the length and width of each tumor using calipers, where V = length × width^2^ × 0.5. Tolerance to intertumoral injections of probiotics was assessed by monitoring mouse weight.

### *In vivo* imaging and biodistribution

All bacterial strains used were luminescent (integrated *luxCDABE* cassette) so they could be visualized with the In Vivo Imaging System (IVIS). To confirm bacterial localization, tumors, spleen and liver were weighed and homogenized using a gentleMACS tissue dissociator (Miltenyi Biotec; C-tubes). Homogenates were serially diluted, plated on LB agar plates and incubated overnight at 37°C. For plasmid retention analysis, tumor homogenates were also plated on LB-agar plates containing kanamycin. Colonies were counted and computed as CFU/g of tissue (limit of detection 10^3^ CFU/g).

### T cell functional assays

To assess cytotoxic responses to D117C and PIGF *ff*Luc^+^ adherent cell lines (HEK293T, HUH7, MCF7, and MDA-MB-68) were allowed to adhere to collagen-coated plates (Thermo Scientific, Nunc™ F96 MicroWell™ White Polystyrene) overnight. The next day media was removed and replaced with serum-free X-VIVO 15 containing D117C or PIGF SA variants and incubated at 37°C while CAR-T cells and untransduced (UT) controls were prepared in serial half-log dilutions to establish a range of effector to target (E:T) ratios. T cells were added to each well and co-cultured for 16-20 hr before addition of Bright-Glo™ (Promega) lysis buffer and luciferin substrate. Luminescence (RLU) was detected with a Tecan plate reader and specific lysis (%) was determined by normalizing RLU to co-cultures with UT T cells. Fold expansion in response to D117C and PIGF SAs was measured by counting T cells (Countess II, ThermoFisher Scientific) every 2-3 days for 14 days following a single stimulation with 0.1 μg/mL of both SAs on collagen coated plates. To assess *in vitro* cytokine production in response to SAs and/or EcN lysate, T cells were washed of IL-2-supplemented media and stimulated for 24 hr as described above. Cells were centrifuged for 5 min at 500 rcf and supernatants were transferred to v-bottom plates (Corning) to clear remaining cellular debris with a second 5 min spin at 500 rcf and transferred to a clean plate for storage at −80°C until cytokine analysis on the Luminex™ 200 (MilliporeSigma, HCD8MAG-17K).

### Flow cytometry

All T cell assays were performed with rested T cells at day 13-15 post activation and purified monomeric GFP was added to all surface stains to stain GFP-CAR^+^ cells or Myc AF488 (Cell Signaling Technology clone 9B11) to stain CD19-, GPC3-, or ICAM1-CAR^+^ cells in flow cytometry-based assays. To assess CD69 expression in response to His-tag purified sfGFP variants, GFP28z CAR-T cells or Jurkats were plated on collagen-coated plates that were pre-incubated with a half-log dilution series of purified sfGFP variants. 16-20 hr later cells were surface-stained with anti-human CD3 BUV395 (BD clone SK7) and CD69 BV421 (Biolegend clone FN50). To assess activation marker status T cells were stimulated with 0.1 μg/mL of D117C or PIGF on collagen coated plates for 16-20 hr and surface-stained with anti-human CD3 BUV395 (BD clone SK7), CD4 BV785 (Biolegend clone RPA-T4), CD8 BV510 (BD clone RPA-T8), CD69 PE-Cy7 (Biolegend clone FN50), CD25 BV421 (Biolegend clone BC96), and CD107a APC (Biolegend clone H4A3). CD107a antibody was added to cell cultures 5 hr before study-end. To assess intracellular cytokine levels T cells were similarly stimulated in the presence of Brefeldin A (BD GolgiPlug) and surface-stained with CD3 BUV395 (BD clone SK7), CD4 BV785 (Biolegend clone RPA-T4) and CD8 BV510 (BD clone RPA-T8), before intracellular staining with TNFα BV421 (Biolegend clone MAb11), IL-2 PE-Cy7 (Biolegend clone MQ1-17H12), IFNγ APC (Biolegend clone B27). Intracellular staining was achieved using a BD Cytofix/Cytoperm™ fixation/permeabilization kit and following manufacturers instruction. To assess activation in response to EcN lysate T cells were similarly stimulated for 16-20 hr with 0.1 μg/mL of PIGF and with, or without, lysate produced by sonication and added at a final OD_600_ of 1. Cells were surface-stained using the activation marker panel above. Finally, to assess T cell phenotype in response to EcN lysate cells were similarly stimulated and surface stained with CD3 BUV395 (BD clone SK7), CD4 BV785 (Biolegend clone RPA-T4), CD8 BV510 (BD clone RPA-T8), CD45RO APC (Biolegend clone UCHL1), and CD62L BV605 (Biolegend clone DREG-56). All samples were acquired on a BD Fortessa.

### *Ex vivo* tumor processing and immunophenotyping

To characterize the effects of live bacteria on human CAR-T cells, Nalm6 tumors were extracted on day 4 post bacteria treatment (day 2 post T cell treatment) and lymphocytes were isolated from tumor tissue by mechanical homogenization using a gentleMACS™ Dissociator (Miltenyi Biotec) in complete hTcm. Cells were filtered through 70 μm cell strainers and washed in PBS before staining. Cells were stained with Ghost Dye™ Red 780 (Cell Signaling Technology) live/dead stain in PBS for 15 min on ice before washing and re-suspending in 2x human/mouse Fc block in stain buffer (BD) followed by surface-staining with monomeric GFP and anti-human CD45 AF700 (Biolegend clone HI30), CD19 PerCP-Cy5.5 (Biolegend clone SJ25C1), CD3 BUV395 (BD clone SK7), CD4 BV785 (Biolegend clone RPA-T4), CD8 BV510 (BD clone RPA-T8), CD45RO APC (Biolegend clone UCHL1), CD62L BV605 (Biolegend clone DREG-56), CD69 PE-Cy7 (Biolegend clone FN50), and CD25 BV421 (Biolegend clone BC96). To characterize T cell exhaustion on day 27 post treatment of MDA-MB-468, T cells were similarly isolated and stained with Ghost Dye and lineage/differentiation markers (without CD19) and with TIM-3 BV711 (Biolegend clone F38-2E2), LAG-3 BV421 (Biolegend clone 11C3C65), PD-1 PE-Cy7 (Biolegend clone EH12.2H7).

To assess *in vivo* production of cytokines in response to EcN, single cell homogenates were similarly achieved from Nalm6 tumors by mechanical homogenization in Milliplex lysis buffer (MilliporeSigma) and Halt™ protease inhibitor (ThermoFisher) and centrifuged at 500 rcf for 10 min to clear debris. Homogenates were transferred to clean tubes for storage at −80°C until cytokine analysis on the Luminex™ 200 (MilliporeSigma, HSTCMAG28SPMX13). sfGFP levels were assessed from similarly prepared tumor homogenates and matched serum drawn from terminal cardiac-puncture using a GFP ELISA kit following manufacturers instruction (abcam).

### Statistical Analysis

Statistical tests were calculated in GraphPad Prism 7.0. The details of the statistical tests are indicated in the respective figure legends. Where data was assumed to be normally distributed, values were compared using a one-way ANOVA for single variable or a two-way ANOVA for more than one variable with the appropriate post-test applied for multiple comparisons. For Kaplan-Meier survival experiments, we performed a log-rank (Mantel-Cox) test.

## Acknowledgments

We thank Dr. Michel Sadelain for kindly providing the *ff*Luc^+^ Nalm6 acute lymphoblastic leukemia cell line, and Dr. Robert Schwabe for kindly providing the Huh7 hepatoma cell line. We thank N. Zervoudis and members of the Obermeyer group for expert assistance with protein purification. We thank M.J. Williams, T. Harimoto, and Dr. M. Rouanne for critical review of the manuscript. Research reported in this publication was performed in the Columbia University Department of Microbiology and Immunology Flow Cytometry core facility and the Irving Institute’s Biomarkers Core Laboratory.

## Author contributions

R.L.V. and T.D. conceived the study. R.L.V. designed and constructed the system, and performed *in vitro* experiments. R.L.V., C.R.G., A.R., and C.C. designed and/or performed *in vivo* experiments. R.L.V., C.R.G., and A.R. performed *ex vivo* experiments. R.L.V., C.R.G., N.A., and T.D. wrote and/or revised the manuscript with input from all authors.

## Data and materials availability

All data is available in the main text or supplementary materials, no datasets were generated or analyzed in the current study. Correspondence and request for materials should be addressed to T.D.

