## supplemental materials for "Probiotic-guided CAR-T cells for universal solid tumor targeting"

### SUPPLEMENTARY MATERIALS

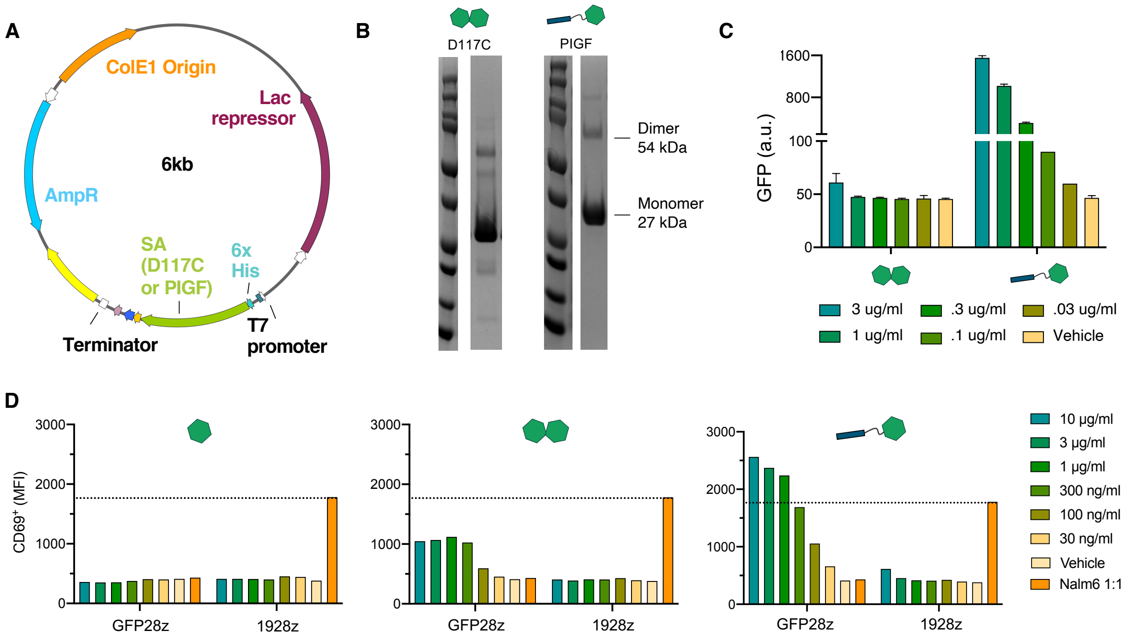


**Supplementary Figure 1. Recombinantly produced, His-tag purified PIGF binds to collagen coated plates and strongly activates GFP-CAR$^+$ Jurkats.** (**A**) Plasmid map of protein expression vector transformed into eNiCo21(DE3) E. *coli* cells for protein purification of D117C and PIGF sfGFP variants. (**B**) Elutions of His-tag purified sfGFP variants, sizes show monomeric and dimeric molecules. (**C**) D117C and PIGF variants were plated in half log dilutions in PBS on collagen coated plates, incubated for 30 m at 37°C and washed 2X with PBS to dislodge any unbound protein. Fluorescence intensity was read at 488 nm on a standard Tecan plate reader. (**D**) CD69 expression on GFP CAR$^+$ or CD19 CAR$^+$ Jurkat cells. Jurkats were plated on collagen coated plates and assessed for CD69 expression by flow cytometry following 16-hour incubation with purified GFP antigens or co-culture with CD19$^+$ target cells (Nalm6). Error bars represent s.d. of biological replicates.

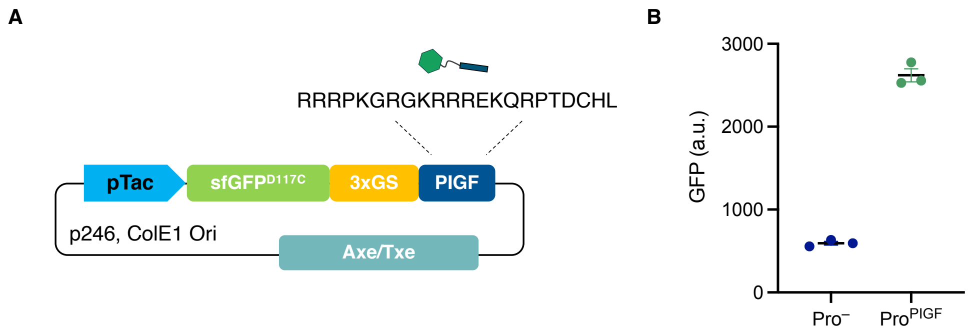


**Supplementary Figure 2. Therapeutic *E. coli* Nissle 1917 (EcN) produces sfGFP-PIGF SA.** (**A**) PIGF SA variant is expressed from an Axe/Txe stabilized, high copy number plasmid under a constitutive *tac* promoter. (**B**) PIGF production by EcN ($Pro^{PIGF}$) as measured by GFP-fluorescence on a Tecan plate-reader and shown relative to an empty EcN control ($Pro^{-}$). Error bars represent s.d. of biological replicates. a.u. arbitrary units.

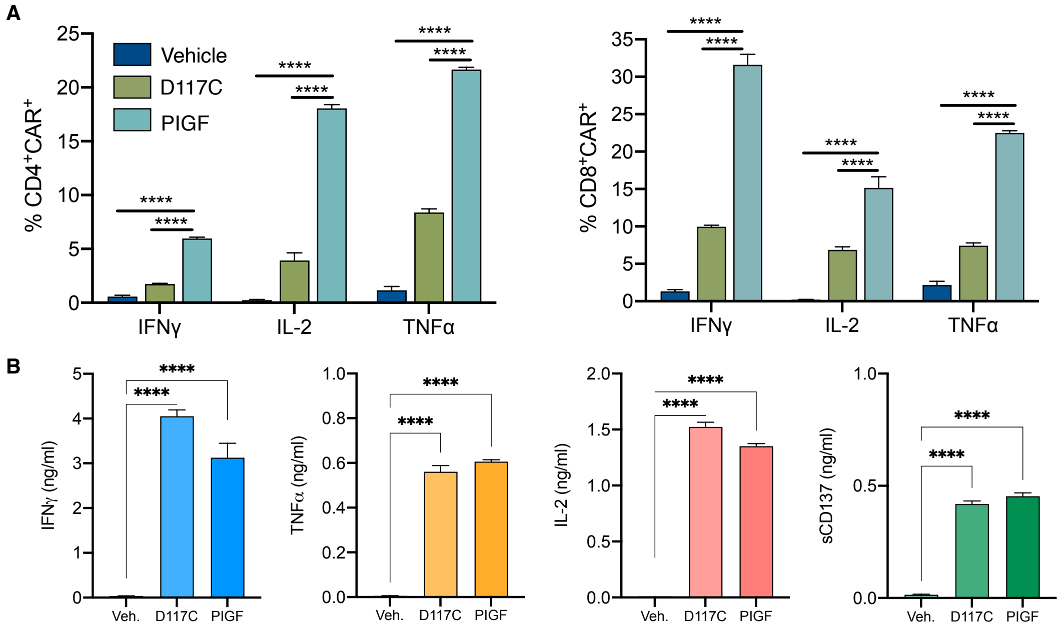


**Supplementary Figure 3. GFP-directed CAR-T cells activate in response to soluble and collagen-bound sfGFP.** (**A**) Quantification of flow cytometric analysis of intracellular staining for pro-inflammatory cytokines in response to 0.1 ng/ml soluble D117C, or collagen bound PlGF. (**B**) Quantification of cytokine production as measured in cell culture supernatants from GFP28z exposed to a PBS vehicle, D117C, or collagen-bound PlGF for 24hr. Error bars represent s.d. of biological replicates, **** $p<0.0001$ 2way (A) or 1way (B) ANOVA, Holm-Sidak multiple comparison correction.

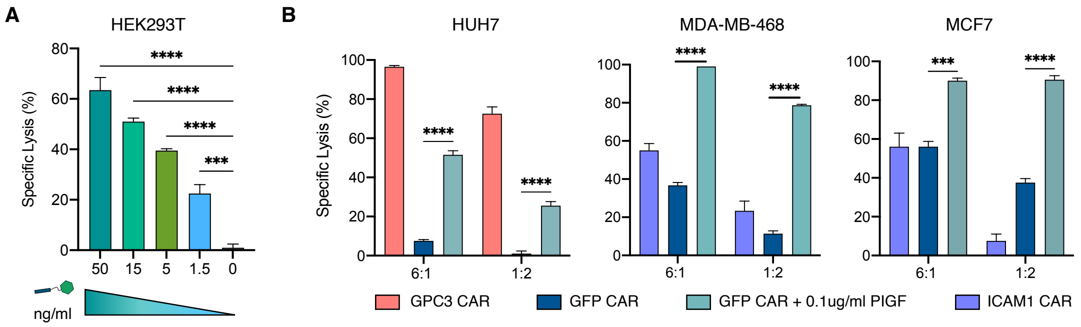


**Supplementary Figure 4. GFP-directed CAR-T cells mediate killing of target cells in response to collagen-bound sfGFP.** (**A**) Overnight killing assays against *ff*Luc$^{+}$ HEK293T at a fixed effector to target (E:T) ratio of 1:3 and half log dilutions of collagen-bound PIGF from 50-1.5 ng/ml. (**B**) Overnight killing assays against *ff*Luc$^{+}$ HUH7, MDA-MB-468, or MCF7 target cells at defined effector to target (E:T). CAR-T cells were co-cultured with target cells on collagen coated plates +/- 0.1 ug/ml sfGFP-PIGF for 20 hours before lysis and addition of luciferin. Luminescence (RLU) was detected with a Tecan plate reader within 20 minutes of lysis. Specific lysis (%) was determined by normalizing RLU to co-cultures with untransduced T cells. Error bars represent s.d. of biological replicates, *** $p<0.001$,**** $p<0.0001$ 1way (A) or 2way ANOVA (B), Holm-Sidak multiple comparison correction.

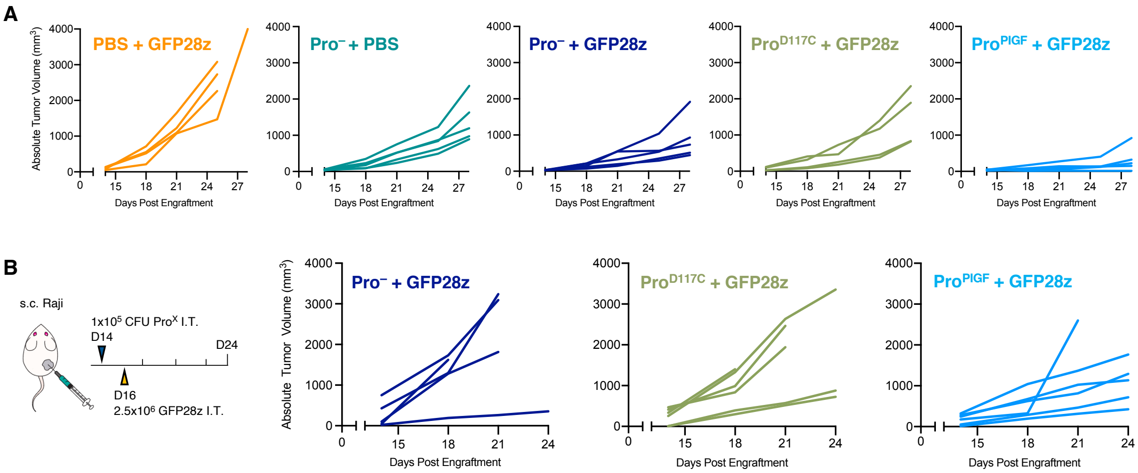


**Supplementary Figure 5. Individual growth trajectories of human tumors treated with the ProCAR system.** (**A**) Nalm6 tumors were established in NSG mice and treated as in Fig. 3A. Individual tumor trajectories are shown. (**B**) Raji lymphoma cells ($5x10^5$) were implanted subcutaneously into the hind flank of NSG mice. When tumor volumes reached ~100 $mm^3$, mice were intratumorally (I.T.) injected with $1x10^5$ CFU of $Pro^-$, $Pro^{D117C}$, or $Pro^{PIGF}$ strains. $2.5x10^6$ GFP28z ProCAR-T cells were then I.T. delivered 48-hours post bacterial injection. Tumor growth was monitored by caliper measurements every 3-4 days, individual tumor trajectories are shown. CFU, colony forming units

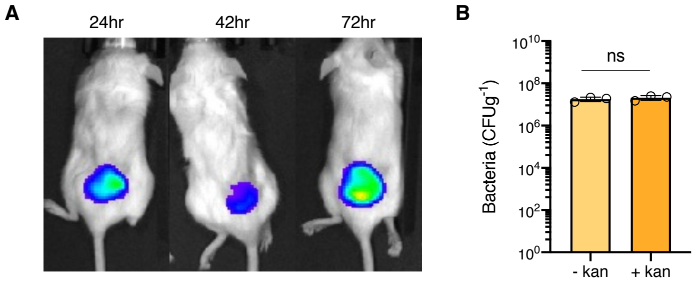


**Supplementary Figure 6. Probiotic EcN remains localized to tumors in immunocompromised NSG mice**. (**A**) IVIS images showing bioluminescent Pro$^-$ populations over time following intratumoral injection of Raji tumors subcutaneously established as in Fig. S5B. (**B**) At day 14 post treatment, Pro$^{PlGF}$-treated tumors were homogenized and plated on LB agar plates containing the appropriate antibiotics (+/- kanamycin) for bacteria colony quantification. Error bars represent s.d. of biological replicates, student's t test; ns, not significant

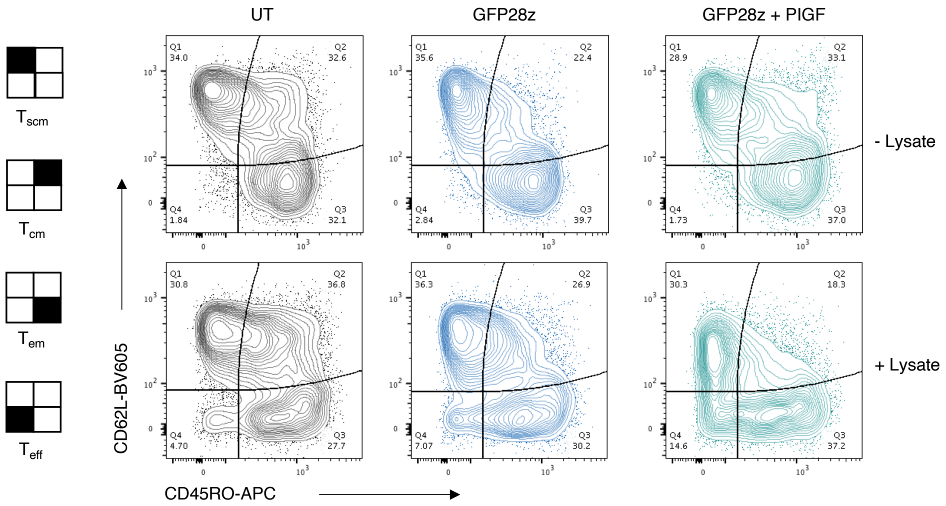


**Supplementary Figure 7. *E. Coli* Nissle (EcN) lysate drives T cell effector phenotype.** Representative flow cytometry contor plots assessing the phenotype of T-cells following stimulation with either media alone or EcN lysate +/- 0.1 ug/ml PIGF for 48hr. $CD8^{+}$ T cell populations were stained for CD45RO and CD62L expression to determine effector T cell differentiation, from stem cell memory ($T_{scm}$) $CD62L^{+}$CD45RO, central memory ($T_{cm}$) $CD62L^{+}CD45RO^{+}$, effector memory ($T_{em}$) $CD62L^{-}CD45RO^{+}$, and terminal effector ($T_{eff}$) $CD62L^{-}$ $CD45RO^{-}$

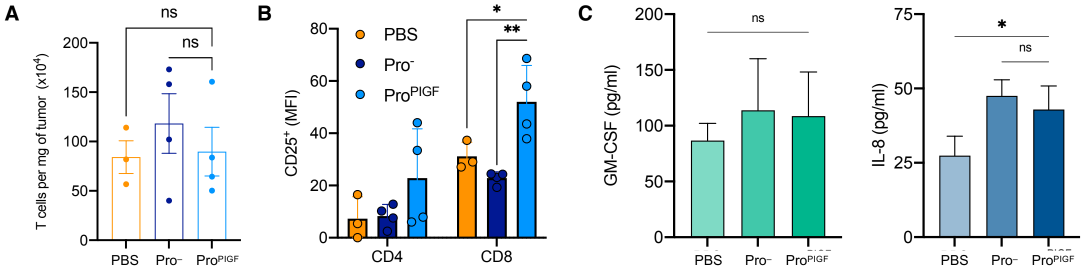


**Supplementary Figure 8. Characterizing the effect of EcN strains on GFP28z cells *in vivo*.** Nalm6 cells ($5x10^5$) were implanted subcutaneously into the hind flank of NSG mice. When tumor volumes reached ~$100mm^3$, mice were I.T. injected with either PBS or $1x10^5$ CFU of $Pro^{PlGF}$ or $Pro^{-}$. On day 2 post $Pro^X$ injection, all groups received an I.T. injection of $2.5x10^6$ GFP28z ProCAR-T cells. Tumors were harvested and homogenized on day 4 for analysis. (**A**) Absolute counts of $hCD45^+CD3^+$ cells per mg of tumor. (**B**) Flow cytometric quantification of CD25 surface expression on intratumoral $hCD45^+CD3^+$ $CD8^+$ or $CD4^+$ cells from each treatment group. (**C**) Quantification of cytokine levels from tumor homogenates. Error bars represent s.d. of biological replicates, * $p<0.05$,** $p<0.01$ 1way (A) or 2way ANOVA (B), Holm-Sidak multiple comparison correction.

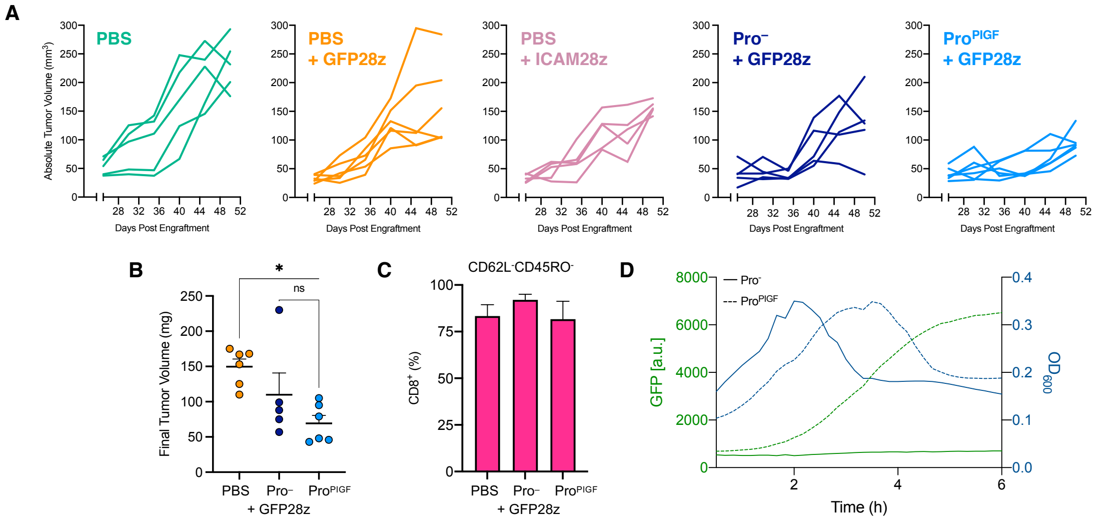


**Supplementary Figure 9. Characterization of T cell exhaustion in a triple negative breast cancer (TNBC) model.** (**A**) Subcutaneous MDA-MB-468 tumors were established in NSG mice prior to I.T. injection with PBS, Pro⁻, or Pro$^{PlGF}$ on day 26 post tumor engraftment. On day 28, mice received a single I.T. injection of either PBS, 2.5x$10^6$ GFP28z ProCAR-T cells, or 2.5x$10^6$ ICAM1-specific CAR-T cells (1CAM28z), as in Fig. 5A. Tumor growth was monitored by caliper measurements every 3-4 days, individual tumor trajectories are shown. (**B**) On day 55 tumors were taken from mice treated with PBS, Pro⁻, or Pro$^{PlGF}$ strains in combination with GFP28z and weighed *ex vivo.* (**C**) Frequency of intratumoral hCD45$^+$CD3$^+$CD8$^+$ cells displaying a terminally differentiated effector phenotype ($T_{eff}$, CD62L$^-$CD45RO$^-$). (**D**) Pro$^X$ strains were isolated from day 55 tumor homogenates and grown overnight on the plate reader to measure $OD_{600}$ and GFP fluorescence intensity. Error bars represent s.d. of biological replicates, * $p<0.05$ 1way ANOVA; ns, not significant

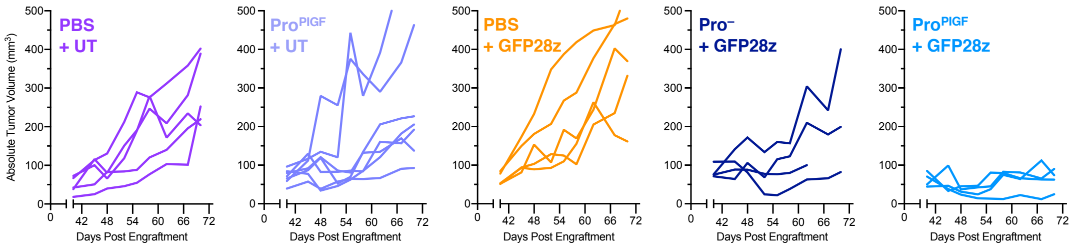


**Supplementary Figure 10. Individual growth trajectories of MDA-MB-468 subcutaneous TNBC tumors treated with the ProCAR system.** Subcutaneous MDA-MB-468 tumors were established in NSG mice prior to I.T. injection with PBS, $Pro^{-}$, or $Pro^{PlGF}$ on day 40 post tumor engraftment. On day 44 Mice received an initial I.T. injection of $2.5x10^6$ untransduced (UT), or $2.5x10^6$ GFP28z ProCAR-T cells, followed by a second I.T. dose of UT or ProCAR-T cells 14 days later (day 58), as in Fig. 5F. Tumor growth was monitored by caliper measurements every 3-4 days, individual tumor trajectories are shown.
